# A CD109-SMURF2 Axis Diverts EGFR from Degradation to Sustain Oncogenic Signaling and Promote Squamous Cell Carcinoma Invasion and Stemness

**DOI:** 10.64898/2026.09.02.748398

**Authors:** Tenzin Kungyal, Amani Hassan, Kenneth Finnson, Varsha Reddy Durgempudi, Anie Philip

## Abstract

Squamous cell carcinoma (SCC) remains difficult to treat, particularly in recurrent or metastatic disease. Although EGFR signaling is a well-established driver of SCC pathogenesis, mechanisms regulating its activity remain incompletely understood. CD109, a GPI-anchored glycoprotein frequently overexpressed in SCC, has been implicated in EGFR signaling, but its mechanisms remain poorly defined. Here, we show that CD109 is elevated in head and neck SCC (HNSCC) in the TCGA PanCancer Atlas and significantly correlates with reduced disease-free survival. Integrative transcriptomic (TCGA PanCancer Atlas) and proteomic (LinkedOmics) analyses revealed strong positive correlations between CD109, EGFR, and SMURF2. These findings were further validated by immunohistochemistry in HNSCC patient samples. Mechanistically, CD109 overexpression impaired EGFR degradation, suppressed degradation-associated EGFR pY1045 phosphorylation, and enhanced signaling-associated pY1068 phosphorylation, resulting in increased STAT3, AKT, and ERK signaling. We identify SMURF2 as a mediator of these effects, with CD109 promoting EGFR-SMURF2 complex formation. SMURF2 gain- and loss-of-function studies in SCC cells, demonstrated that SMURF2 stabilizes EGFR by reducing EGFR interaction with the ubiquitin ligase c-Cbl, thereby limiting EGFR turnover in a CD109-dependent manner. Functionally, the CD109-SMURF2-EGFR axis promotes EGF-induced stemness, invasion, and proliferation in SCC cells. Together, these findings identify CD109 as a molecular switch controlling EGFR fate by redirecting EGFR from c-Cbl-mediated degradation toward SMURF2- dependent stabilization, sustaining oncogenic signaling. This previously unrecognized regulatory pathway reveals the CD109-SMURF2 axis as a vulnerability with potential for targeted intervention in EGFR-driven SCCs.

## 1. Introduction

Squamous cell carcinoma (SCC), arising from squamous cells of the skin and mucous membranes, is a leading cause of cancer-related mortality worldwide [1,2]. Oral SCC (OSCC) accounts for over 90% of oral malignancies in North America, with rising incidence driven by tobacco, alcohol, and HPV infection [3]. Five-year survival ranges from roughly 50-65% depending on site and stage, with better outcomes in HPV-positive disease [4]. While early-stage lesions respond well to surgery or radiation, advanced or recurrent disease often demands more aggressive treatment [5], underscoring the need to delineate the molecular mechanisms driving progression to identify new therapeutic targets [6].

Epidermal growth factor receptor (EGFR, ErbB1/HER1) is a transmembrane receptor tyrosine kinase that drives cell growth and differentiation upon EGF binding [7–10] and plays a central role in SCC pathogenesis, particularly in head and neck SCC (HNSCC), including OSCC [11]. EGFR is overexpressed in approximately 80-90% of HNSCC cases, correlating with poor prognosis, tumor aggressiveness, and therapeutic resistance [12]. Unlike lung adenocarcinoma, where activating mutations such as L858R or exon 19 deletions predominate, EGFR dysregulation in SCC arises primarily through amplification and overexpression rather than mutation [12,13]. The monoclonal antibody cetuximab, an FDA-approved EGFR-targeted therapy for HNSCC, is often used alongside radiotherapy or chemotherapy [14], yet therapeutic resistance remains a persistent challenge, motivating continued research into alternative inhibitors, combination strategies, and predictive biomarkers [14].

CD109 is a glycosylphosphatidylinositol (GPI)-anchored protein and a member of the α2- macroglobulin (α2M)/C3, C4, C5 complement family of thioester-containing proteins [19–21]. Our group first defined CD109’s functional role, showing that it acts as a transforming growth factor-β (TGF-β) co-receptor that negatively regulates TGF-β1 signaling and thereby restrains downstream responses such as extracellular matrix synthesis and fibrosis [19,22–25], in part by promoting SMURF2-dependent degradation of the TGF-β receptor [26,27]. Consistent with this tumor-suppressive function, CD109 potently inhibits TGF-β-induced epithelial-mesenchymal transition (EMT) in A431 SCC cells [1]. Rather than simply blocking EMT, CD109 appears to promote a terminally differentiated epithelial state, thereby restraining the hybrid epithelial/mesenchymal phenotype required for migration and metastasis [2]. CD109 expression is nonetheless elevated in SCC across multiple tissues - including skin, oral cavity, lung, vulva, and bladder [30–38] – and premalignant SCC lesions with high CD109 expression carry an increased risk of metastatic progression [33], implicating CD109 in cancer progression. Furthermore, our group [15] and others [16] have shown that CD109 deletion *in vivo* blocks cancer progression by decreasing EGFR signaling in SCC, a mechanism also reported in lung adenocarcinoma [17] and glioblastoma [18].

EGFR activity is tightly controlled not only by ligand-induced phosphorylation but also by the trafficking route the receptor follows activation. Upon EGF binding, EGFR is rapidly internalized into early endosomes, from which it is either recycled back to the plasma membrane or routed to lysosomes for degradation - a critical step that determines the duration and intensity of downstream signaling [3]. EGFR degradation is governed principally by ubiquitination, catalyzed by the E3 ubiquitin ligase c-Cbl. Efficient c-Cbl recruitment depends on phosphorylation of EGFR at Y1045, its direct c-Cbl-binding site, together with indirect, GRB2-dependent recruitment via Y1068 and Y1086 [4]; loss of Y1045 phosphorylation impairs c-Cbl association and receptor ubiquitination, resulting in reduced degradation and enhanced EGFR recycling to the cell surface [5, 6]. Y1068 and Y1086 also recruit adaptors such as Grb2 or Gab1 leading to activation of STAT3, AKT, and ERK signaling [7, 8], and the balance between c-Cbl-dependent degradation and receptor recycling directly shapes the amplitude and duration of oncogenic EGFR signaling[9, 10]. Thus, in addition to receptor abundance or ligand availability, EGFR’s oncogenic output in cancer cells, is modulated by c-Cbl-EGFR engagement.

Here we show that high CD109 expression correlates with increased EGFR expression and poor disease-free survival in SCC. CD109 enhances EGF-mediated STAT3, AKT, and ERK signaling in SCC (A431, FaDu) cells, increasing EGF-induced EGFR phosphorylation at Y1068 - the Grb2/Gab1 docking site driving downstream signaling - while decreasing phosphorylation at Y1045, the c-Cbl docking site required for receptor ubiquitination and degradation. Consistent with this, CD109 overexpression abrogated EGF-dependent EGFR-c-Cbl interaction and reduced EGF-induced EGFR ubiquitination in A431 cells. Bioinformatic analysis of HNSCC samples further showed that CD109 expression positively correlates with SMURF2, an E3 ligase that, unlike most others, stabilizes rather than degrades EGFR [41]. Importantly, CD109 promoted EGFR protein stability and EGF-induced EGFR-SMURF2 interaction and colocalization, as well as EGF-induced stem cell marker expression (Oct4, Nanog, Sox2) and invasion, across SCC cell lines and primary HNSCC cells, in a CD109-dependent manner. Together, these findings support a model in which CD109 stabilizes EGFR and sustains its signaling both by promoting SMURF2- EGFR interaction and by suppressing c-Cbl-mediated EGFR ubiquitination and degradation - identifying CD109 as a novel regulator of EGFR fate that links receptor trafficking to oncogenic signaling in SCC.

## 2. Materials and methods

### 2.1 cBioPortal and LinkedOmics database analysis

The cBioPortal for Cancer Genomics serves as a publicly available online platform for the interactive exploration of cancer genomics datasets (http://www.cbioportal.org/) [11, 12]. CD109 mRNA expression data (RNA Seq V2) were obtained from 32 cancer studies via the cBioPortal interface. The cBioPortal database was queried using the keywords CD109, EGFR, and Smurf2 to analyze expression correlations within cohorts from The Cancer Genome Atlas (TCGA) PanCancer Atlas, including HNSCC, Cervical SCC, and Lung SCC (n = 523). Furthermore, the HNSCC cohort was analyzed to assess survival outcomes in patients with high and low CD109 expression levels. For the correlation study between CD109 and EGFR, processed proteomics data files were retrieved from LinkedOmics (n=109 patients) https://www.linkedomics.org/login.php [13]. The patients were categorized into two groups based on CD109 expression levels: the top 25% for high CD109 expression and the bottom 25% for low CD109 expression. Pearson correlation analysis between CD109 and EGFR protein levels in these patient groups was performed using GraphPad Prism.

### 2.2 Microarray and Gene set enrichment analysis (GSEA)

The microarray analysis has been previously published and described elsewhere [1]. All genes, irrespective of their differential expression status, were utilized for GSEA. GSEA was conducted using GSEA software (version 4.1.0) [14] to examine gene sets exhibiting statistically significant differences between the microarray data of CD109 wild type (WT) and CD109 KO A431 cells. We selected oncogenic gene sets from the gene set database and calculated normalized enrichment scores (NES) and adjusted q-values using the GSEA method, which was based on 1000 random permutations of the ranked gene list.

Collection of tumor and adjacent normal tissue specimens from patients with oral HNSCC samples were obtained from the Department of Otorhinolaryngology – Head and Neck Surgery at McGill University. The study received approval from the Research Ethics Committee of the Jewish General Hospital (Protocol #11-093) and was conducted in accordance with established guidelines. Written informed consent was obtained from all participants, and experimental protocols were approved by the Canadian Research Review Office (RRO). Eligible patients were HPV-negative, had no prior treatment, and did not present with a second primary tumor, with all seeking care at the same institution. Human HNSCC cells were isolated from the tongue tumor of patients as described elsewhere [15].

### 2.3 Cell culture

The human vulvar squamous carcinoma cell line A431 (CRL-1555) (ATCC, Manassas, VA, USA) was cultured in Dulbecco’s modified Eagle’s medium (DMEM, Gibco, Thermo Fisher Scientific, USA) supplemented with 10% fetal bovine serum (FBS) and 1% penicillin-streptomycin (PS). The human HNSCC cell line FaDu (HTB-43) (ATCC, Manassas, VA, USA) was maintained in minimum essential medium (MEM, Gibco, Thermo Fisher Scientific, USA) supplemented with 10% FBS and 1% PS. A431 CD109 knockout (KO) cells were generated using CRISPR/Cas9 genome editing, as described in our previous study [16]. FaDu CD109 KO cells, targeted with a CRISPR/Cas9 gRNA (CUUCCAGGACAGAGACAGUG) at exon 2, were obtained from Synthego (USA). CD109 overexpressing (OE) and empty vector (EV) A431 stable cell lines were cultured in DMEM supplemented with 10% FBS, 1% PS, and geneticin (Gibco; #10131-035). FaDu cell populations and primary human HNSCC cells with high and low CD109 expression were sorted by fluorescence-activated cell sorting (FACS).

### 2.4 Western blot

Cell lysates (20 μg of protein) were subjected to immunoblotting as previously described[17]. In brief, the cell extracts were separated using SDS-polyacrylamide gel electrophoresis and transferred to nitrocellulose membranes. The membranes were blocked with blocking buffer (5% BSA in TBST) at room temperature for 45 minutes, and subsequently incubated overnight at 4°C with primary antibodies for CD109 (Santa Cruz, sc271085), EGFR (Santa Cruz, sc-373746), pEGFR 1068 (Santa Cruz, sc-377547), pEGFR 1045 (Santa Cruz, sc-57541), STAT3 (Santa Cruz, sc-8019), pSTAT3 (Santa Cruz, sc-8059), AKT (Cell Signaling Technology, 4691S), pAKT (Santa Cruz, sc-514032), pERK (Santa Cruz, sc-7383), ERK (Cell Signaling, 4695), Smurf2 (G-Biosciences, ITA7689), and c-Cbl (Santa Cruz, sc-1651). Following incubation, the membranes were washed three times with TBST and then probed with secondary antibodies (Cell Signaling, 7074S or 7076S) for 1 hour at room temperature. Protein bands were visualized using Clarity™ Western ECL Substrate (Bio-Rad).

### 2.5 Co-immunoprecipitation (Co-IP) Study

Co-IP assays were performed as described previously [18]. Briefly, cells were lysed with Pierce IP lysis buffer (87797), and the lysates were mixed with Protein G magnetic beads (Bio-Rad, 1614023) preincubated with primary antibody for 1hr at room temperature on a rotator shaker. Beads were washed with PBST three times to remove unbound proteins. The bead bound proteins were eluted with 2X Laemmli buffer for 10 min at 70°C followed by immunoblotting.

### 2.6 Transfection

The siRNA transfection protocol was conducted as previously described [19]. siRNA targeting Smurf2 or control scrambled siRNA (Thermo Fisher Scientific, 4392420, S34859) were transfected into cells using Lipofectamine™ RNAiMAX Transfection Reagent (Thermo Fisher Scientific, Waltham, MA, USA, 13778030). At 35 hours post-transfection, cells were serum-starved overnight and subsequently treated with EGF as specified.

The CD109 pcDNA3.1 plasmid, pHAGE EGFR (116731), and pCMV5B-Flag-Smurf2 (11746), along with their corresponding EVs, were transfected into A431 CD109 KO cells using Lipofectamine 2000 Transfection Reagent (Thermo Fisher Scientific, 11668030) following the manufacturer’s instructions.

### 2.7 Immunohistochemistry

For patient tumor immunohistochemistry (IHC) analysis, the protocol was followed as previously described [20]. Briefly, tissue sections were treated with 3% hydrogen peroxide for 5 minutes at room temperature to block endogenous peroxidase activity. This was followed by blocking with 0.1% BSA in PBS for 1 hour. The sections were then incubated with the primary antibody overnight at 4°C. Subsequent incubation with a biotinylated secondary antibody was carried out for 30 minutes, followed by incubation with streptavidin for 1 hour at room temperature. The slides were developed using a DAB substrate solution and counterstained with hematoxylin.

### 2.8 MTT Assay

Colorimetric MTT (3-(4,5-dimethylthiazol-2-yl)-2,5-diphenyl tetrazolium bromide, 475989 Sigma) assays were performed to assess the mitochondrial metabolic activity, an indirect measure of cell viability [21]. Following transfection, media was removed and replaced with serum-free media containing MTT (0.5 mg/mL) and cells were then incubated for four hours at 37° C. The formazan formed was dissolved with dimethyl-sulfoxide (DMSO, Sigma) and absorbance was measured at 595 nm using a microplate reader.

### 2.9 Statistical analysis

Statistical analyses were performed using multiple comparative methods: Student’s t-test for paired comparisons, one-way ANOVA with post-hoc analysis for multiple group comparisons, two-way ANOVA for examining the effects of two independent variables, and Chi-square test for categorical data analysis. All experimental procedures were conducted in biological triplicates to ensure reproducibility. Data are expressed as mean ± SEM and were analyzed using GraphPad Prism software (v9.0). Statistical significance was established at p < 0.05 for all analysis. * p < 0.05, ** p < 0.01, *** p < 0.001, **** p < 0.0001

## 3. Results

### 3.1 High CD109 expression correlates with poor disease-free survival and elevated EGFR levels in SCC patients

To define the clinical relevance of CD109 in squamous malignancies, we interrogated The Cancer Genome Atlas (TCGA) PanCancer Atlas datasets through cBioPortal. Across 32 tumor types, CD109 mRNA expression was most prominent in SCCs, with the highest levels observed HNSCC, followed by lung and cervical SCCs (Fig. 1A, Supplementary FigS1). These results demonstrate that CD109 expression is selectively enriched in SCC tumors. We next examined the prognostic significance of CD109 expression in HNSCC. Kaplan-Meier analysis revealed that tumors with high CD109 expression were associated with significantly reduced disease-free survival compared with CD109-low tumors (Fig. 1B), supporting a role for CD109 expression in disease progression. Given our prior microarray data implicating CD109 in regulation of EGFR signaling [2], we decided to evaluate the relationship between CD109 and EGFR expression in clinical specimens. Analysis of TCGA transcriptomic data demonstrated a strong positive correlation between CD109 and EGFR mRNA levels in HNSCC (Fig. 1C). Consistent with this finding, interrogation of LinkedOmics proteomic datasets confirmed a positive association between CD109 and EGFR protein expression in HNSCC specimens (Fig. 1D). Together, these data indicate that CD109 expression correlates with elevated EGFR mRNA and protein levels and adverse clinical outcome in HNSCC, providing clinical relevance for the mechanistic studies described below.

**Figure 1.**
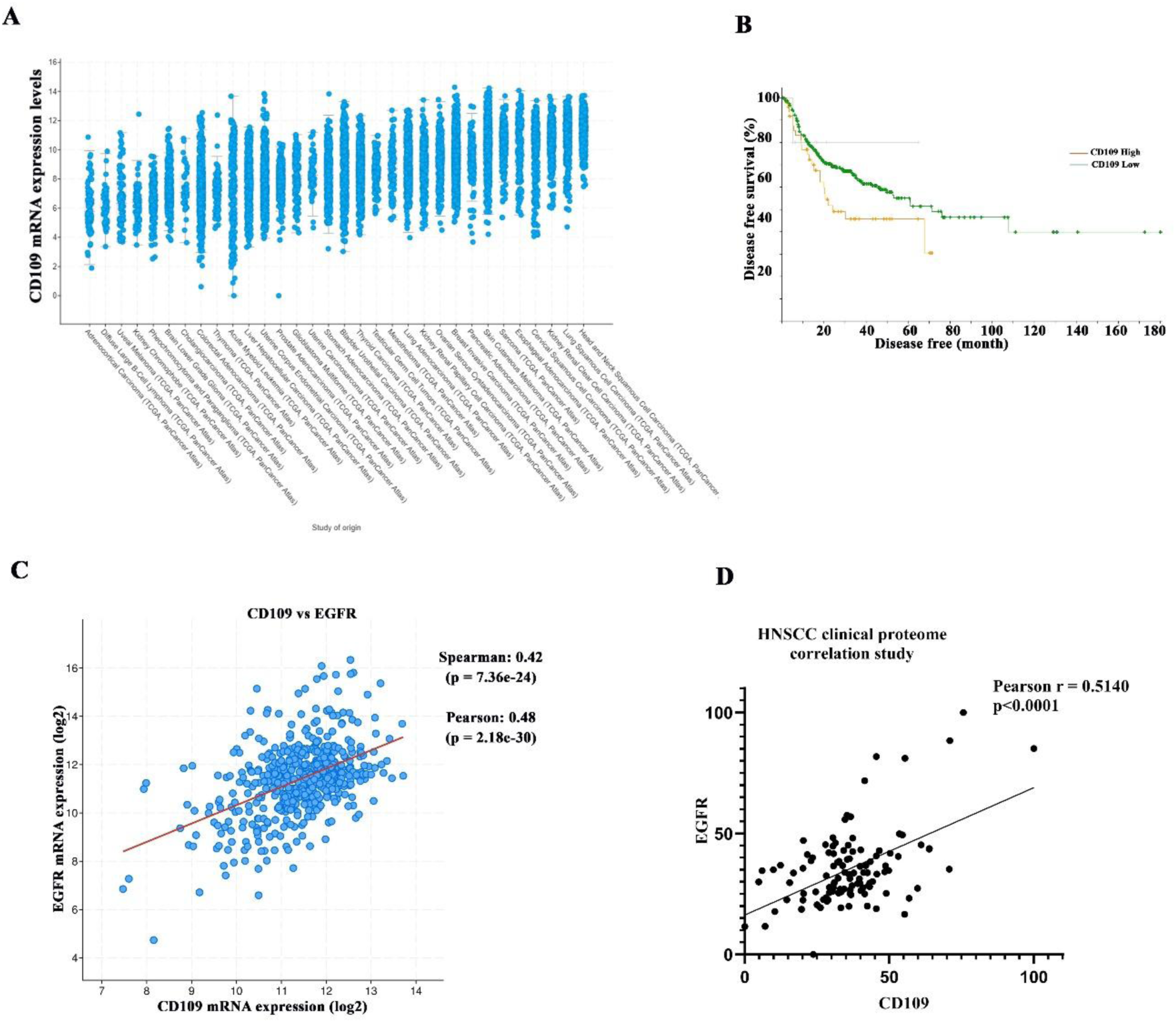
High CD109 expression correlates with poor disease-free survival and elevated EGFR levels in SCC patients. **(A)** Pan-cancer analysis of CD109 mRNA expression across multiple tumor types using The Cancer Genome Atlas (TCGA) datasets accessed through the cBioPortal platform. **(B)** Kaplan-Meier analysis of disease-free survival in head and neck squamous cell carcinoma (HNSCC) patients stratified by CD109 expression levels (high vs. low) based on TCGA cohort data. Survival differences were evaluated using the log-rank test. **(C)** Correlation analysis of CD109 and EGFR mRNA expression in HNSCC patients (n = 523) from TCGA. Pearson correlation coefficient (r) and corresponding P value are indicated. **(D)** Clinical proteomic analysis demonstrating correlation between CD109 and EGFR protein expression levels in HNSCC specimens (LinkedOmics database). Statistical significance is defined as *P < 0.05.

### 3.2 CD109 attenuates EGFR degradation and promotes EGFR stability and signaling in SCC cells

To interrogate the contribution of CD109 to oncogenic signaling, we performed gene set enrichment analysis (GSEA) of microarray profiles from CD109 KO and WT A431 cells. We found that EGFR-associated oncogenic signatures were significantly enriched in CD109 WT relative to CD109 KO A431 cells (Fig. 2A) suggesting that CD109 expression correlates with EGFR mRNA levels in SCC cells. This is consistent with our previous report that CD109 enhances EGFR mRNA levels in SCC cells, as determined by quantitative PCR [2]. We next examined CD109 regulates EGFR signaling using WT vs CD109 KO A431 and FaDu SCC cells. In both models, CD109 KO cells displayed decreased EGF-induced EGFR phosphorylation and decreased activation of downstream effectors, including STAT3, AKT, and ERK, compared with CD109 WT cells (Fig. 2B, C). Notably, CD109 deficiency was also associated with reduced total EGFR protein levels, suggesting that CD109 regulates both EGFR abundance and EGFR signaling output.

**Figure 2.**
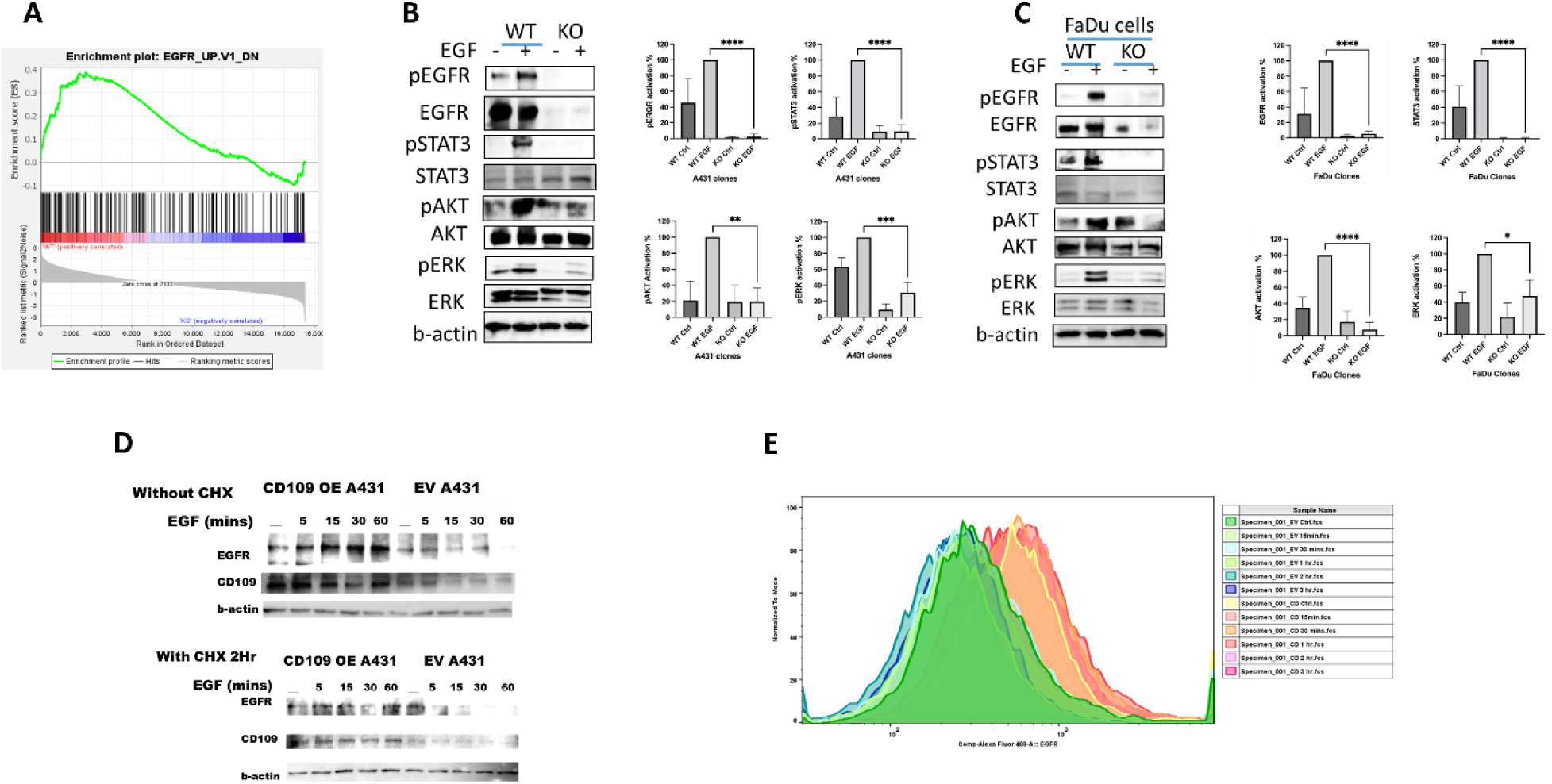
CD109 attenuates EGFR degradation and promotes EGFR stability and signaling in SCC cells. **(A)** Gene Set Enrichment Analysis (GSEA) of microarray data comparing A431 wild-type (WT) and CD109 knockout (CD109-KO) cells, demonstrates attenuation of the EGFR oncogenic signature in CD109-KO cells. Normalized enrichment score (NES) and false discovery rate (FDR) are indicated. **(B)** Western blot analysis of EGFR and downstream signaling pathway activation in A431 WT and CD109-KO cells following EGF stimulation (100 ng/mL) for the indicated time points. **(C)** Western blot analysis of EGFR signaling in FaDu WT and CD109-KO cells following EGF treatment. **(D)** EGFR protein stability assessed by cycloheximide (CHX) chase assay in A431 empty vector (EV) and CD109-overexpressing (CD109-OE) cells. Cells were treated with EGF alone (upper panel) or EGF plus CHX (lower panel) for the indicated time points, followed by Western blot analysis of EGFR. Quantification of EGFR levels normalized to time 0 is shown. **(E)** Flow cytometric analysis of cell surface EGFR expression in A431 EV and CD109- OE cells. Representative histograms and quantification of mean fluorescence intensity (MFI) are presented. Data represent mean ± SEM from at least three independent experiments. Statistical significance was determined using unpaired two-tailed Student’s t-test or two-way ANOVA with appropriate post hoc testing. *P < 0.05; **P < 0.01; ***P < 0.001.

To determine whether CD109 influences EGFR protein stability, we performed cycloheximide (to block de novo protein synthesis) chase assays to assess EGFR degradation . While EGFR levels declined following EGF stimulation in empty vector transfected (EV) cells, EGFR degradation was markedly delayed in CD109-overexpressing (OE) A431 cells (Fig. 2D), consistent with enhanced EGFR stability. Consistent with this, flow cytometric analysis revealed that CD109- overexpressing cells retained markedly higher surface EGFR levels than EV controls following EGF treatment—an effect evident as early as 15–30 minutes, timepoints at which de novo protein synthesis is negligible, implicating a role for CD109 in decreased EGFR trafficking rather than its synthesis (Fig. 2E). Collectively, these data indicate that CD109 sustains EGFR signaling in SCC cells by decreasing EGFR degradation and thereby promoting EGFR stability.

### 3.3 CD109 inhibits EGF-induced EGFR ubiquitination and EGFR-c-Cbl interaction in SCC cells

To investigate the role of CD109 in regulating EGFR stability, we compared CD109 OE A431 cells with EV controls following treatment with EGF for varying time points. Analysis revealed a marked reduction in EGFR ubiquitination in CD109 OE cells, as illustrated (Fig. 3A, left panel). This reduction was further supported by a significant decrease in phosphorylation of EGFR at tyrosine 1045, a site associated with the receptor’s degradation pathway (Fig. 3A, middle panel) [22]. These findings suggest that EGF-induced EGFR degradation is diminished in the presence of CD109 overexpression. Conversely, we observed a significant increase in EGFR phosphorylation at tyrosine 1068, indicative of enhanced downstream signaling in these CD109 OE cells. In addition, CD109 overexpression led to inhibition of EGF-induced EGFR interaction with c-Cbl (responsible for EGFR degradation), thereby preventing EGFR degradation (Fig. 3B). On the other hand, EGF treatment promoted the interaction of c-Cbl with EGFR in WT A431 cells (Supplementary fig. S2).. Collectively, these data demonstrate that CD109 plays a critical role in maintaining EGFR stability and enhancing its signaling potential.

**Figure 3.**
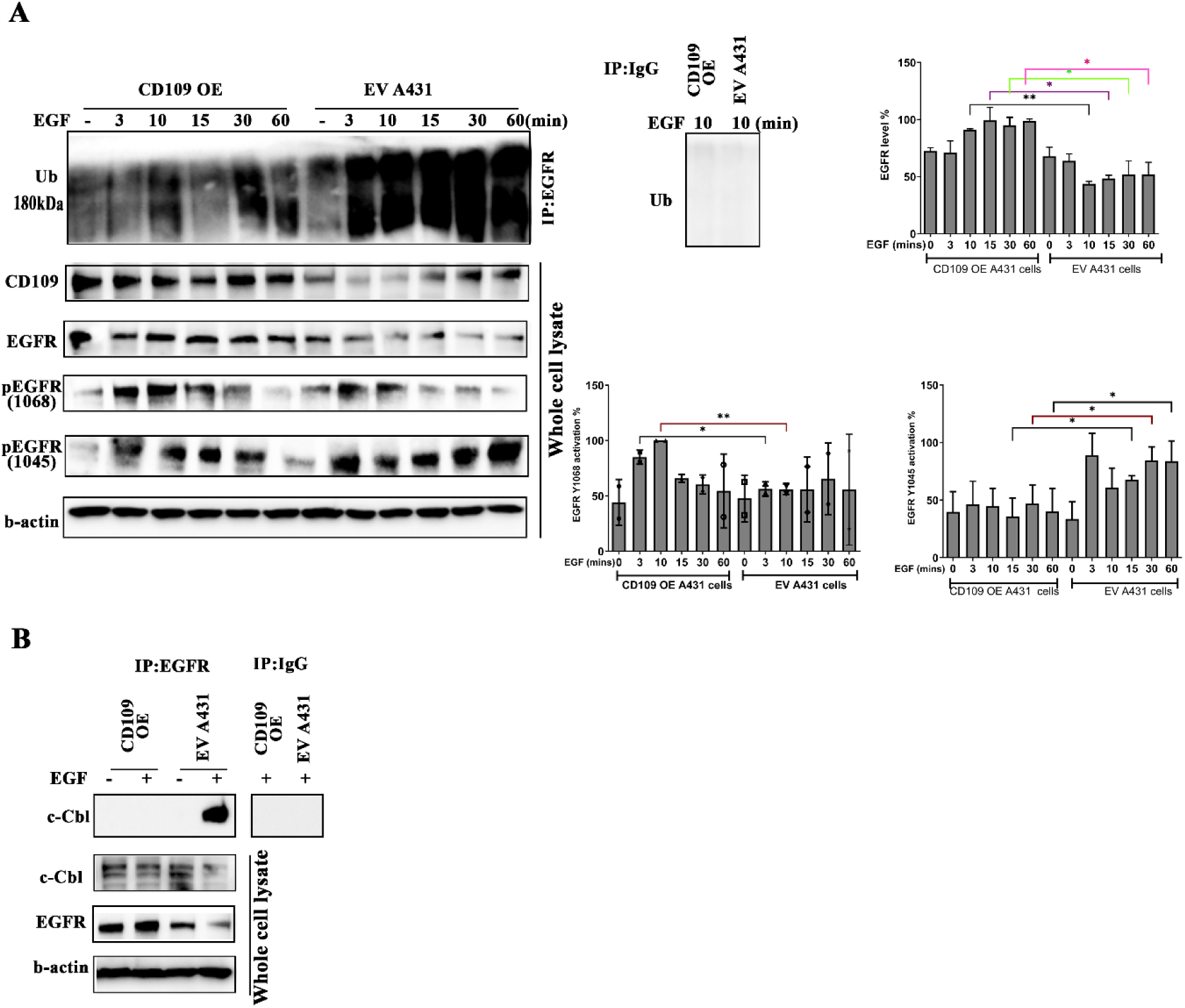
CD109 inhibits EGF-induced EGFR ubiquitination and EGFR-c-Cbl interaction in SCC cells. **(A)** A431 cells stably expressing empty vector (EV) or CD109-overexpression (CD109-OE) were stimulated with EGF (100 ng/ml) for the indicated time points. EGFR ubiquitination was assessed by immunoprecipitation of EGFR followed by immunoblotting for ubiquitin. Total EGFR levels and phosphorylation at Y1045 and Y1068 were analyzed by Western blotting. Densitometric quantification normalized to loading control or total EGFR is shown. **(B)** Co-immunoprecipitation analysis of EGFR-c-Cbl interaction in A431 EV and CD109-OE cells following EGF stimulation. EGFR was immunoprecipitated and probed for c-Cbl by Western blotting. Quantification of EGFR-bound c-Cbl relative to total EGFR is presented. Data represent mean ± SEM from at least three independent experiments. Statistical significance was determined using unpaired two-tailed Student’s t-test. *P < 0.05; **P < 0.01; ***P < 0.001.

### 3.4 CD109-dependent stabilization of EGFR is mediated by Smurf2

Smurf2 has been previously shown to stabilize EGFR by antagonizing c-Cbl-mediated ubiquitination [23]. We therefore examined whether CD109 modulates EGFR stability through a Smurf2-mediated mechanism. Immunohistological analysis showed a positive correlation of CD109 with EGFR and Smurf2 in HNSCC patient tissues (Supplementary fig. S3). Analysis of our previous microarray data [2] revealed a significant reduction in Smurf2 mRNA levels in CD109 KO A431 cells compared to WT (Fig. 4A). Supporting this, TCGA data from HNSCC patients show a positive correlation between CD109 and Smurf2 expression (Fig. 4B). We then examined Smurf2 protein expression patterns using CD109 loss or gain of function (OE and KO SCC cells) and showed that CD109 enhances Smurf2 expression (Fig. 4C). We next investigated the importance of Smurf2 expression on EGFR stability. Our results indicate that siRNA-mediated knockdown of Smurf2 results in a marked decrease in EGFR protein levels (Fig. 4D). To investigate the interactions among CD109, EGFR, and Smurf2, as well as their individual effects, we transiently overexpressed these proteins either alone or in various combinations in CD109-KO A431 cells. Our findings demonstrated that EGFR expression was highest when all three proteins - CD109, EGFR, and Smurf2 - were co-expressed, compared with their expression levels when each protein was expressed alone or in other combinations (Fig. 4E). Collectively, these data indicate that CD109 and Smurf2 act in concert to augment EGFR stability, consistent with a synergistic regulatory mechanism.

**Figure 4.**
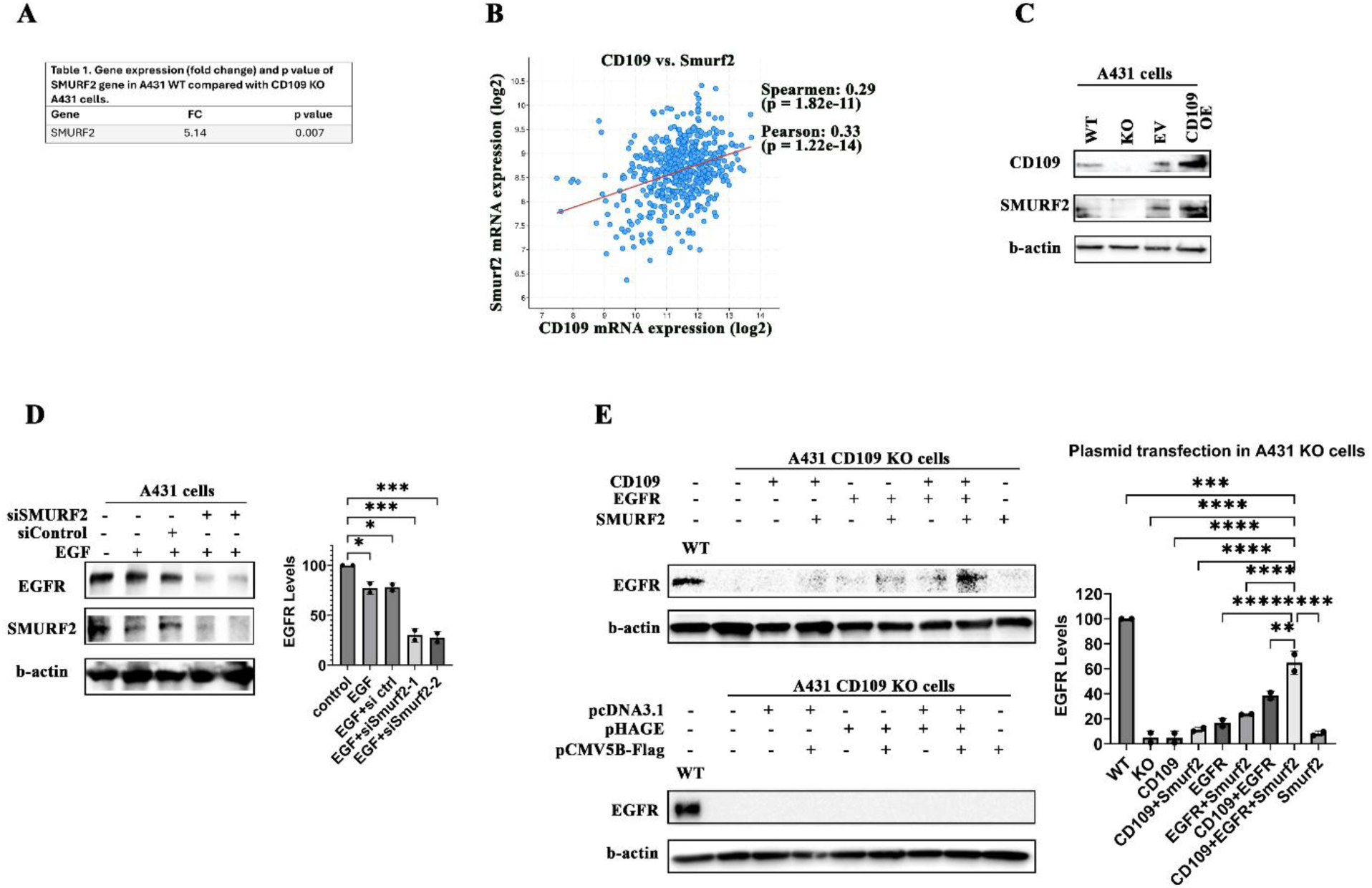
CD109-dependent stabilization of EGFR is mediated by Smurf2. **(A)** Microarray analysis comparing Smurf2 mRNA expression in A431 wild-type (WT) and CD109 knockout (CD109-KO) cells. Differential expression values are shown. **(B)** Correlation analysis of CD109 and Smurf2 mRNA expression in head and neck squamous cell carcinoma (HNSCC) patient samples (n = 523) from The Cancer Genome Atlas (TCGA) dataset accessed via the cBioPortal platform. Pearson correlation coefficient (r) and P value are indicated. **(C)** Western blot analysis of Smurf2 protein expression in A431 cells with varying CD109 levels (WT, CD109-KO, empty vector [EV], and CD109-overexpression [CD109-OE]). Densitometric quantification normalized to loading control is shown. **(D)** Effect of Smurf2 knockdown on EGFR protein levels in A431 cells. Cells were transfected with control or Smurf2 siRNA, and EGFR expression was analyzed by Western blotting. Quantification of EGFR levels relative to control is shown. **(E)** CD109-KO A431 cells were transiently transfected with expression plasmids encoding CD109, Smurf2, EGFR, or the indicated combinations, along with corresponding empty vector controls. EGFR protein levels were analyzed by Western blotting (upper and lower panels as indicated). Quantification of EGFR expression normalized to loading control is presented. The data represent mean ± SEM from at least three independent experiments. Statistical significance was determined using unpaired two-tailed Student’s t-test or one-way ANOVA with appropriate post hoc testing. *P < 0.05; **P < 0.01; ***P < 0.001.

### 3.5 CD109 enhances Smurf2-mediated EGFR stability by inhibiting EGFR-c-Cbl interaction in SCC cells

To elucidate the mechanism underlying the requirement of Smurf2 for CD109-mediated EGFR stabilization, we depleted Smurf2 in A431 WT cells using siRNA and subsequently assessed alterations in the EGFR-c-Cbl interaction by co-immunoprecipitation. Smurf2 depletion enhanced EGFR-c-Cbl interaction, which was accompanied by a reduction in total EGFR protein levels (Fig. 5A). We extended these findings to primary HNSCC cells, which were stratified into CD109-high and CD109-low populations by flow cytometry. CD109-high primary HNSCC cells exhibited diminished EGFR-c-Cbl interaction, and a concomitant increase in total EGFR protein levels. Smurf2 siRNA knockdown abrogated this effect, restoring EGFR-c-Cbl interaction and reducing EGFR stability (Fig. 5B). In addition, CD109-high primary HNSCC cells display an increase in EGF-induced phosphorylation of EGFR at Y1068 (activation) and decreased EGF-induced phosphorylation at Y1045 (degradation) (Fig. 5B) consistent with an increase in EGFR signaling and decrease in EGFR degradation.

**Figure 5.**
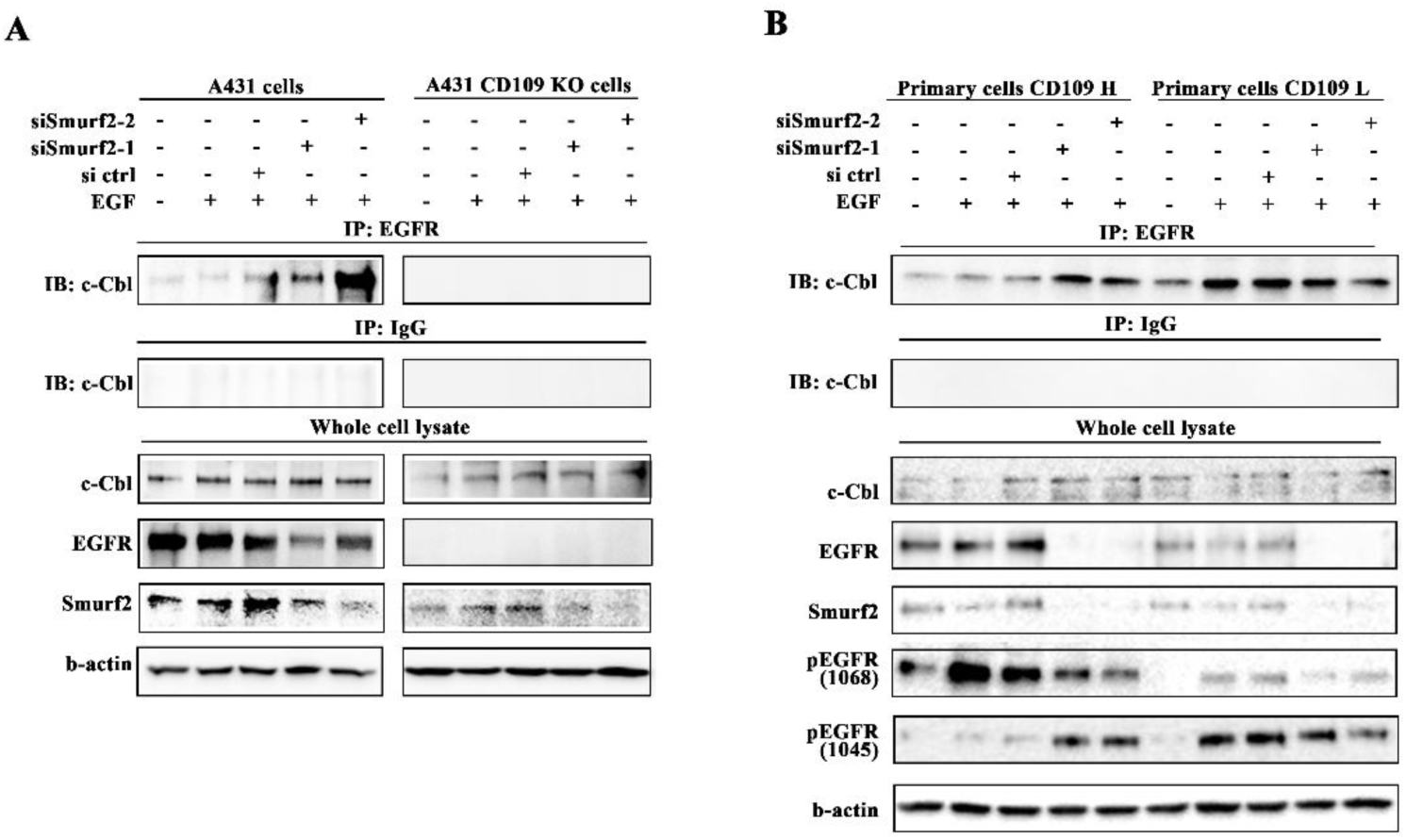
CD109 enhances Smurf2-mediated EGFR stability by inhibiting EGFR-c-Cbl interaction in SCC cells. **(A)** A431 wild-type (WT) and CD109 knockout (CD109-KO) cells were transfected with control or Smurf2 siRNA and stimulated with EGF (100 ng/mL) for the indicated time points. EGFR-c-Cbl interaction was assessed by co-immunoprecipitation. EGFR was immunoprecipitated and probed for c-Cbl by Western blotting. Quantification of EGFR-bound c- Cbl relative to total EGFR is shown. **(B)** A431 empty vector (EV) and CD109-overexpressing (CD109-OE) cells were subjected to Smurf2 knockdown followed by EGF stimulation. EGFR-c- Cbl interaction was analyzed by co-immunoprecipitation as described above. Representative immunoblots and densitometric quantification are presented. The data represent mean ± SEM from at least three independent experiments. Statistical significance was determined using unpaired two- tailed Student’s t-test or one-way ANOVA with appropriate post hoc testing. *P < 0.05; **P < 0.01; ***P < 0.001.

### 3.6 CD109 promotes EGFR stability by enhancing EGFR-Smurf2 association in SCC cells

Endocytic trafficking plays a crucial role in regulating EGFR signaling and degradation [24]. To determine whether CD109 influences EGFR cellular distribution or association with Smurf2, we assessed total EGFR levels, cell surface EGFR abundance, and EGFR-Smurf2 colocalization by immunofluorescence confocal microscopy. CD109-overexpressing (OE) A431 cells exhibited significantly higher total EGFR protein levels and sustained cell surface EGFR following EGF stimulation compared with EV cells (Fig. 6A). Similarly, CD109-high primary HNSCC cells maintained elevated total and cell surface EGFR levels relative to CD109-low counterparts (Fig. 6B). Furthermore, CD109 OE A431 cells and CD109-high primary cells revealed enhanced colocalization of EGFR with Smurf2. Thus, CD109 may stabilize EGFR by promoting its association with Smurf2, thereby limiting c-Cbl recruitment, reducing degradative ubiquitination, and favoring EGFR stability and sustained oncogenic signaling.

**Figure 6.**
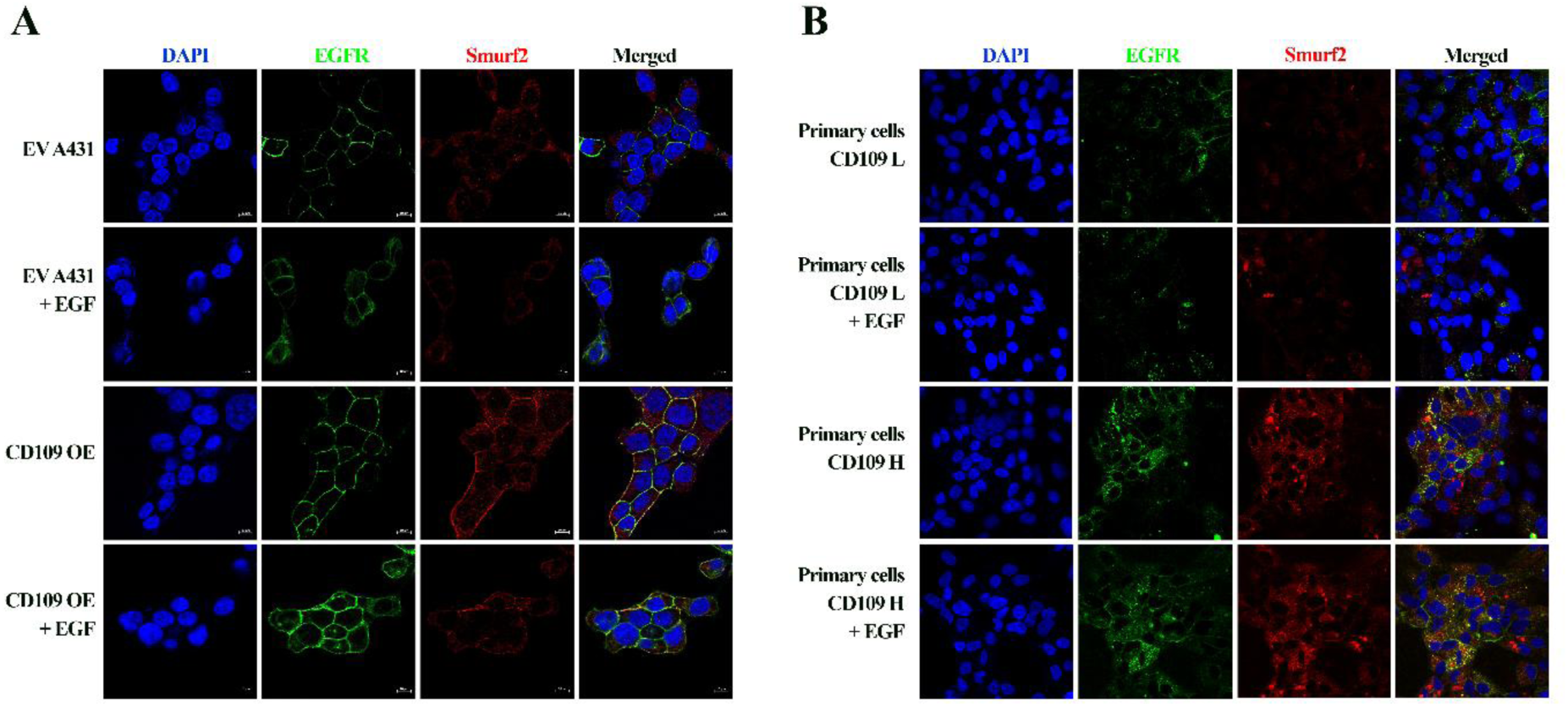
CD109 promotes EGFR stability by enhancing EGFR-Smurf2 association in SCC cells. **(A)** A431 cells stably expressing empty vector (EV) or CD109-overexpression (CD109-OE) were stimulated with EGF (100 ng/mL) for the indicated time points. EGFR-Smurf2 colocalization was assessed by immunofluorescence confocal microscopy. **(B)** CD109-indcued EGFR-Smurf2 colocalization was examined in primary head and neck squamous cell carcinoma (HNSCC) cells stratified by CD109 expression (CD109-high vs. CD109-low). Representative images from at least three independent experiments are shown.

### 3.7 CD109 promotes EGFR-driven stem cell marker expression via Smurf2

To determine whether CD109/EGFR-driven stemness is mediated by Smurf2, we used A431 and FaDu SCC cells with CD109 overexpression or knockout in the presence or absence of EGF. EGF stimulation induced expression of the stem cell transcription factors Oct4, Nanog, and Sox2, with maximal induction observed at 6 h (Fig. 7A, B). Our results show that siRNA-mediated Smurf2 depletion led to attenuated stemness marker expression (Oct4, Nanog and Sox2) in KO cells, whereas WT cells showed no significant reduction of those markers.(Fig. 7C). Consistent with these findings, CD109-OE cells displayed enhanced EGF-induced stem cell marker expression. Importantly, this increase was significantly diminished following Smurf2 silencing (Fig. 7D).

**Figure 7.**
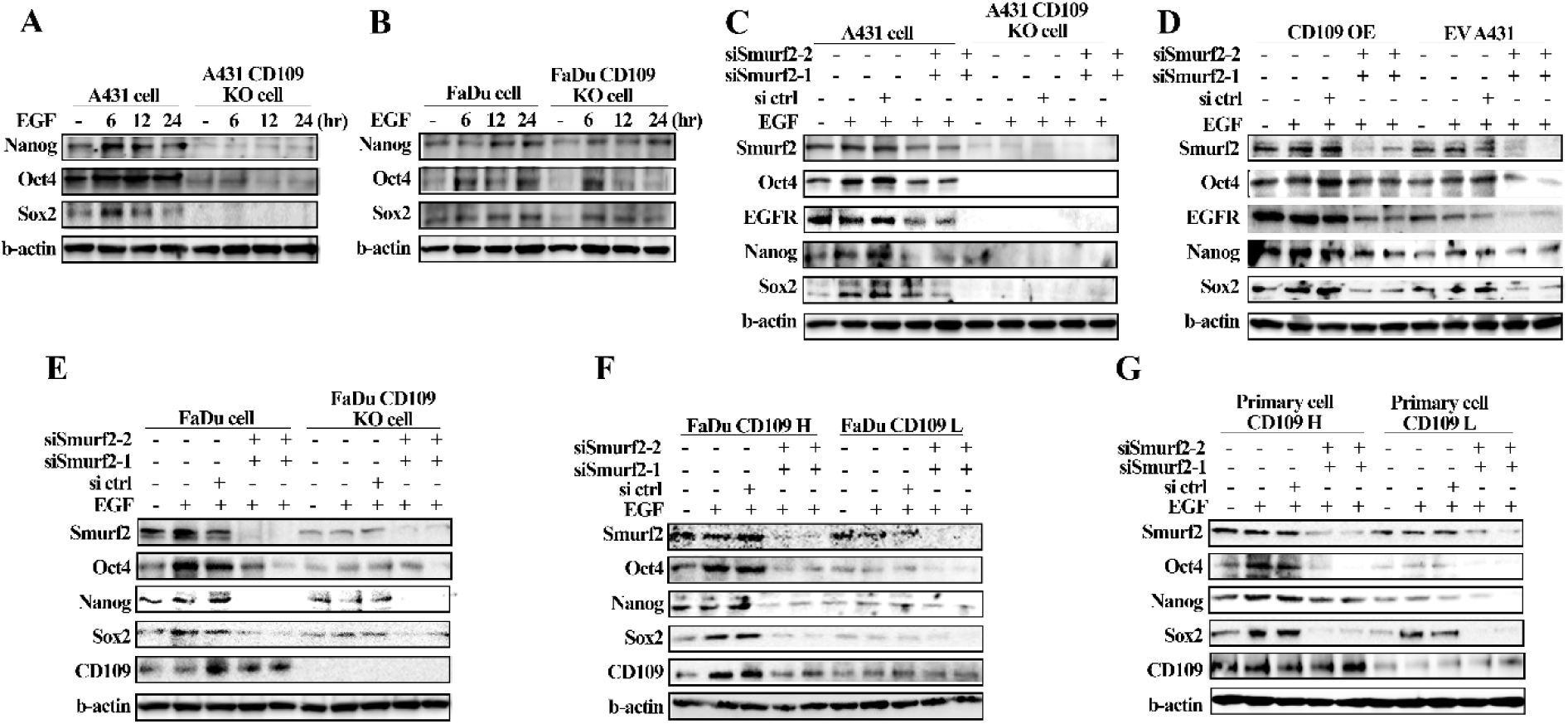
CD109 promotes EGFR-driven stem cell marker expression via Smurf2. **(A, B)** Time-course analysis of stem cell marker expression following EGF stimulation (100 ng/mL) in A431 (A) and FaDu (B) cells. Wild-type (WT) and CD109 knockout (CD109-KO) cells were treated with EGF for the indicated time points, and the expression of Oct4, Nanog, and Sox2 was analyzed by Western blotting. Representative Western blots and densitometric quantification normalized to loading control are shown. **(C)** A431 WT and CD109-KO cells were transfected with control or Smurf2 siRNA and stimulated with EGF. Expression of Oct4, Nanog, and Sox2 was analyzed by Western blotting. **(D)** A431 empty vector (EV) and CD109-overexpressing (CD109-OE) cells were subjected to Smurf2 knockdown followed by EGF stimulation, and stem cell marker expression was assessed as described above. **(E)** FaDu WT and CD109-KO cells were transfected with control or Smurf2 siRNA and treated with EGF prior to analysis of Oct4, Nanog, and Sox2 expression. **(F)** FaDu cells sorted into CD109-high and CD109-low populations were transfected with control or Smurf2 siRNA, stimulated with EGF, and analyzed for stem cell marker expression by Western blotting. **(G)** Primary head and neck squamous cell carcinoma (HNSCC) cells stratified by CD109 expression (CD109-high vs. CD109-low) were subjected to Smurf2 knockdown followed by EGF stimulation, and Oct4, Nanog, and Sox2 expression levels were analyzed by Western blotting. Data represent mean ± SEM from at least three independent experiments. Statistical significance was determined using unpaired two-tailed Student’s t-test or one-way/two-way ANOVA with appropriate post hoc testing. *P < 0.05; **P < 0.01; ***P < 0.001.

We next validated these findings in FaDu cells. Smurf2 knockdown in WT and CD109-KO FaDu cells similarly reduced EGF-mediated induction of Oct4, Nanog, and Sox2 (Fig. 7E). Furthermore, in FaDu populations sorted for high or low CD109 expression, CD109-high cells exhibited greater EGF-induced stem cell marker expression, which was markedly attenuated upon Smurf2 depletion (Fig. 7F). Finally, primary HNSCC cells stratified by CD109 expression were analyzed. CD109- high primary cells demonstrated robust EGF-induced stemness marker expression, and this response was significantly reduced following Smurf2 knockdown (Fig. 7G). Densitometric analyses of these data are shown in supplementary figure S4. Taken together, these results demonstrate that Smurf2 is required for CD109-dependent amplification of EGF-induced stemness across multiple SCC cell models.

### 3.8 CD109 promotes EGF-induced invasion and proliferation via Smurf2

In the tumor microenvironment, EGF acts as a chemotactic agent, promoting the motility and invasion of tumor cells, thereby enhancing invasion and metastasis [25]. In this context, we assessed the ability of SCC cells to invade and migrate directionally toward EGF. EGF-induced invasion was markedly increased in SCC cells overexpressing CD109 and in CD109-high primary HNSCC cells (Fig. 8A). This enhanced invasive capacity was substantially reduced following Smurf2 knockdown, indicating a functional requirement for Smurf2 in CD109-dependent, EGF-mediated cell invasion. Consistent with these findings, MTT assays showed that CD109-OE A431 cells exhibited significantly increased cell viability compared with EV A431 controls (Fig. 8B-D). Notably, this CD109-driven increase in viability was completely abolished by Smurf2 knockdown, demonstrating that Smurf2 is essential for CD109/EGFR-dependent regulation of SCC cell proliferation. Together, these findings link CD109-Smurf2 axis to EGFR stabilization and activation resulting in enhanced invasive and proliferative behavior in SCC cells.

**Figure 8.**
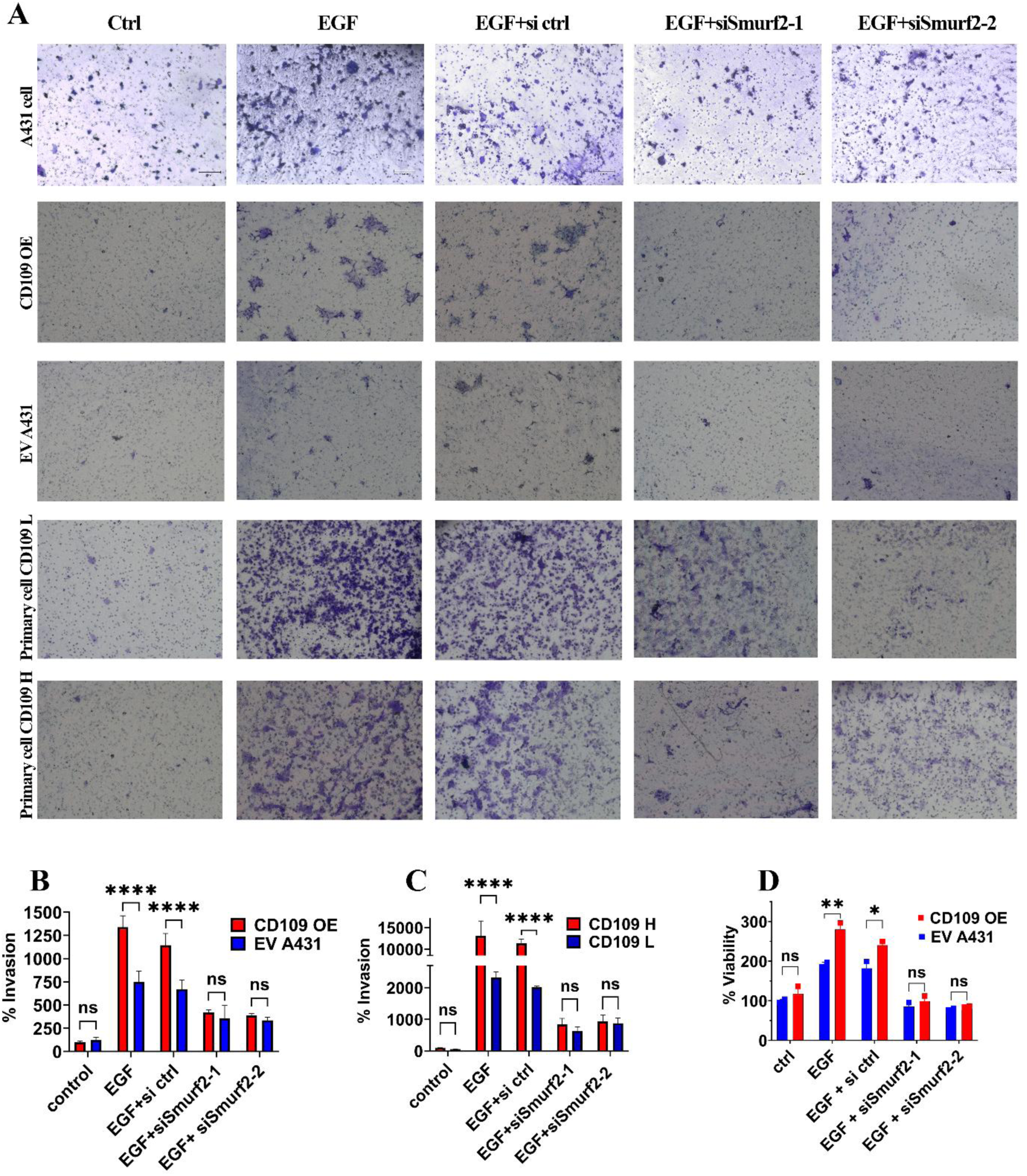
CD109 promotes EGF-induced invasion and proliferation via Smurf2. **(A)** Invasion assays were performed to evaluate the effect of Smurf2 knockdown on EGF-induced invasive capacity. Parental A431 cells, A431 empty vector (EV), CD109-overexpressing (CD109-OE) A431 cells, and primary head and neck squamous cell carcinoma (HNSCC) cells stratified by CD109 expression (CD109-high vs. CD109-low) were transfected with control or Smurf2 siRNA and stimulated with EGF (100 ng/mL). Representative images of invaded cells are shown. **(B, C)** Quantification of invaded cells from the assays shown in **(A)**. Invasion was measured in **(B)** A431 EV and CD109-OE cells and **(C)** primary HNSCC cells stratified by CD109 expression. Results are expressed as the number of invaded cells per field relative to control conditions. **(D)** Cell proliferation was assessed using an MTT assay in A431 EV and CD109-OE cells following Smurf2 knockdown and EGF stimulation. Proliferation kinetics were measured at the indicated time points. Data represent mean ± SEM from at least three independent experiments. Statistical significance was determined using unpaired two-tailed Student’s t-test or one-way/two-way ANOVA with appropriate post hoc testing. *P < 0.05; **P < 0.01; ***P < 0.001.

## Discussion

In this study, we show that CD109 governs the fate of EGFR, acting as a control point that dictates EGFR downstream trafficking or turnover in SCC cells. By acting as a regulatory switch CD109 determines whether EGFR is degraded or retained/recycled by redirecting EGFR away from c-Cbl-mediated degradation through SMURF2 engagement, thus uncovering a previously unrecognized route by which SCC cells sustain oncogenic EGFR signaling. We demonstrate that CD109 promotes Smurf2-dependent stabilization of EGFR, by shifting EGFR away from c-Cbl-mediated degradation to sustain its oncogenic signaling. Mechanistically, CD109 enhances EGFR- Smurf2 interaction, attenuates phosphorylation at the degradative Y1045 site, increases phosphorylation at the signaling-associated Y1068 site, and reduces EGFR ubiquitination and turnover. Analyses of public databases (TCGA and LinkedOmics) indicate that elevated CD109 expression correlates with reduced disease-free survival in head and neck SCC (HNSCC), and increased EGFR and Smurf2 levels, underscoring the potential clinical relevance of this molecular signaling network. These findings define CD109 as a determinant of receptor fate after ligand engagement and establish a mechanistic framework linking CD109 expression to persistent EGFR signaling and tumor progression.

EGFR signaling intensity and duration are tightly controlled by post-activation trafficking of EGFR and its ubiquitination that determine whether EGFR is recycled or degraded [9, 10, 26]. EGF-induced phosphorylation of Y1068 facilitates recruitment of adaptor proteins such as Grb2 and activation of downstream mitogenic pathways [27], whereas phosphorylation at Y1045 promotes recruitment of the E3 ubiquitin ligase c-Cbl, triggering EGFR ubiquitination and lysosomal degradation [5, 6, 28]. Our data shows that CD109 overexpression reduces EGF-induced EGFR degradation, maintains higher cell surface EGFR levels, and maintains activation of STAT3, AKT, and ERK pathways. These effects are accompanied by reduced EGF-induced phosphorylation of EGFR at Y1045 and enhanced phosphorylation at Y1068 suggesting that CD109 influences this decision point. Together, these findings suggest that CD109 shifts EGFR signaling toward continued mitogenic and survival pathway activation by modulating its ubiquitination and trafficking.

A key discovery of this study is that CD109-driven EGFR stabilization is critically Smurf2- dependent. Our results demonstrate that siRNA-mediated Smurf2 knockdown reduces EGFR protein levels, indicating that Smurf2 contributes to EGFR stability. In CD109 KO cells, re-expression of CD109 restores EGFR protein levels, an effect that is further enhanced by Smurf2 overexpression. Consistent with these findings, CD109 promotes EGFR-Smurf2 colocalization, supporting a cooperative role for CD109 and Smurf2 in stabilizing EGFR. Importantly, our results demonstrate that CD109 overexpression reduces EGF-induced EGFR ubiquitination and disrupts EGF-induced EGFR-c-Cbl interaction, suggesting that CD109 protects EGFR from c-Cbl-mediated ubiquitination and subsequent degradation. Furthermore, loss of Smurf2 restores EGFR-c-Cbl interaction, accelerates receptor degradation, and abolishes the enhanced signaling and invasive behavior of CD109-high cells, suggesting that Smurf2 is required for CD109-dependent inhibition of c-Cbl recruitment and the stabilization of EGFR. While Smurf2 has been previously implicated in modulating EGFR stability by antagonizing c-Cbl-mediated ubiquitination [23, 29], our findings reveal CD109 as a key upstream orchestrator that promotes Smurf2 recruitment to EGFR in SCC cells, which disrupts EGFR-c-Cbl interaction to promote EGFR stability. These results reveal a novel CD109-Smurf2-EGFR axis as a critical regulator of EGFR stability and signaling in SCC, highlighting a targetable therapeutic vulnerability to suppress SCC tumor invasiveness.

Sustained EGFR signaling has been linked to cancer stem-like properties, therapeutic resistance, and invasive behavior [27]. Here we show that CD109-high cells display Smurf2-dependent enhancement of EGF-induced stem cell transcription factors (Nanog, Oct4, Sox2), increased proliferation, and greater invasiveness in SCC cells suggesting that EGFR stabilization by CD109- Smurf2 drives functional reprogramming. By stabilizing EGFR, the CD109-Smurf2 axis may sustain downstream signaling pathways that drive stemness-associated gene expression and promote invasive behavior.

Smurf2 is classically recognized as a HECT-domain E3 ubiquitin ligase regulating TGF-β receptor and Smad turnover [30–35]. Beyond its role in TGF-β signaling, Smurf2 has been implicated in modulating EGFR stability and tumor progression [23, 29], and its effects on proliferation and invasion have been shown to be context-dependent [36]. We have previously shown that CD109 negatively regulate TGF-β signaling by promoting receptor internalization and degradation in a Smad7/Smurf2-dependent manner[37, 38]. In this regard, our findings presented here suggest that CD109/Smurf2 exerts opposing effects on distinct receptor systems: facilitating TGF-β receptor degradation while stabilizing EGFR. This dual role of CD109/Smurf2 axis could shift the signaling balance from tumor-suppressive TGF-β pathway toward pro-oncogenic EGFR signaling, a transition frequently observed during SCC progression [39]. Our analysis of public cancer genomics datasets provides important clinical relevance for the functional studies presented here. Interrogation of TCGA PanCancer Atlas data through cBioPortal demonstrated that CD109 expression is markedly enriched in SCCs compared to many other tumor types, with the highest levels observed in HNSCC. Within the HNSCC cohort, patients with high CD109 expression exhibited significantly reduced disease-free survival, supporting a functional role in disease progression and also suggesting that CD109 may serve as a prognostic indicator of aggressive disease. These findings are consistent with previous reports demonstrating that elevated CD109 expression is associated with adverse clinical outcomes, including reduced overall survival, decreased disease-free survival, increased tumor aggressiveness, and enhanced metastatic potential in oral, vulvar and lung SCC[40–42].

To our knowledge, prior studies have not systematically examined the relationship between CD109 and EGFR expression at the transcriptomic and proteomic level in large, publicly available cancer datasets. Here, analysis of TCGA PanCancer Atlas (transcriptomic) and LinkedOmics (proteomic) data sets reveal a strong positive association between CD109 and EGFR expression across both transcript and protein expression. This relationship was further validated at the protein level, by immunohistochemical analysis of HNSCC patient samples, which revealed a significant positive correlation between CD109 and EGFR, as well as between CD109 and Smurf2, supporting the notion that CD109 contributes to enhanced EGFR signaling in human tumors. Additionally, our results show for the first time that CD109 expression positively correlates with Smurf2 levels in HNSCC samples consistent with a coordinated regulatory axis. In this regard, our finding that the expressions of CD109, EGFR, and Smurf2 are all higher in the tumor tissue relative to adjacent normal tissue in HNSCC patients (Supplementary Fig. 3) further supports this inference, underscoring their potential significance in HNSCC pathology.

Mechanistically, our findings support a model in which CD109 facilitates recruitment or stabilization of Smurf2 at activated EGFR complexes. This interaction reduces phosphorylation at Y1045, limiting c-Cbl docking and degradative ubiquitination, while maintaining phosphorylation at Y1068, thereby preserving downstream signaling competence. The net effect is a shift in EGFR fate from lysosomal degradation toward sustained signaling and surface persistence. This mechanism aligns with emerging models in which modulation of ubiquitination dynamics determines receptor signaling duration and oncogenic output [43]. In this framework, CD109 functions not merely as a co-receptor but as a regulator of receptor quality control, biasing ubiquitination-dependent trafficking decisions toward tumor-promoting signaling.

Building on this mechanism, several aspects of the pathway remain to be determined. While we demonstrate that CD109 reduces EGFR ubiquitination, the specific ubiquitin linkage architecture underlying this shift - for example, K48- versus K63-linked chains or monoubiquitination - remains to be defined. Given that distinct ubiquitin topologies dictate divergent trafficking outcomes [74], resolving how CD109 shapes chain architecture represents a natural next step toward a complete mechanistic picture of this regulatory axis. Similarly, although Smurf2 knockdown abolishes CD109-mediated EGFR stabilization, establishing this axis firmly, the precise contribution of Smurf2 catalytic activity versus a scaffolding role remains to be determined; catalytically inactive Smurf2 mutants offer a direct route to this distinction. Finally, an important translational extension of this work will be to determine whether CD109 expression modulates sensitivity to EGFR-targeted therapies, a question of relevance in HNSCC, where EGFR overexpression rather than activating mutation predominates [75].

In conclusion, our study establishes CD109 as a pivotal regulator of EGFR stability and responses in SCC, acting through Smurf2 to bias EGFR fate away from degradative pathways and toward sustained oncogenic signaling. By simultaneously modulating EGFR and TGF-β receptor dynamics, CD109 may orchestrate a broader shift in the signaling landscape that favors tumor progression, stemness, and invasiveness. These findings position the CD109-Smurf2-EGFR axis as a critical driver of EGFR stability and signaling in SCC, and a potential targetable vulnerability and biomarker for aggressive disease. Future studies dissecting the precise ubiquitin architectures involved, the necessity of Smurf2 enzymatic activity, and the impact of CD109 on responsiveness to EGFR-targeted therapies will be critical for translating this mechanistic insight into clinical interventions aimed at overcoming EGFR-driven resistance in HNSCC.

## Data availability

Data are available on request. The remaining data are available within the Article, and Supplementary Information.

## CRediT authorship contribution statement

Tenzin Kungyal: review & editing, Writing – original draft, Validation, Supervision, Project administration, Formal analysis, Visualization, Data curation, Conceptualization. Amani Hassan, Varsha Reddy Durgempudi and Kenneth Finnson: Formal analysis, Data curation, review and editing. Anie Philip: Writing – review & editing, Supervision, Project administration, Funding acquisition, Conceptualization.

## References

[1] S. Zhou, S.D. da Silva, P.M. Siegel, A. Philip, CD109 acts as a gatekeeper of the epithelial trait by suppressing epithelial to mesenchymal transition in squamous cell carcinoma cells in vitro, Scientific reports, 9 (2019) 16317.

[2] S. Zhou, A. Hassan, T. Kungyal, S. Tabaries, J. Luna, P.M. Siegel, A. Philip, CD109 is a critical determinant of EGFR expression and signaling, and tumorigenicity in squamous cell carcinoma cells, Cancers, 14 (2022) 3672–3785.

[3] G. Jendrisek, D. Mesa, A. Conte, M.G. Malabarba, S. Sigismund, Beyond the membrane: rethinking EGFR signaling in physiology and cancer, Cellular and molecular life sciences : CMLS, 83 (2026) 109.

[4] F. Capuani, A. Conte, E. Argenzio, L. Marchetti, C. Priami, S. Polo, P.P. Di Fiore, S. Sigismund, A. Ciliberto, Quantitative analysis reveals how EGFR activation and downregulation are coupled in normal but not in cancer cells, Nature communications, 6 (2015) 7999.

[5] H. Waterman, G. Levkowitz, I. Alroy, Y. Yarden, The RING finger of c-Cbl mediates desensitization of the epidermal growth factor receptor, The Journal of biological chemistry, 274 (1999) 22151–22154.

[6] G. Levkowitz, H. Waterman, E. Zamir, Z. Kam, S. Oved, W.Y. Langdon, L. Beguinot, B. Geiger, Y. Yarden, c-Cbl/Sli-1 regulates endocytic sorting and ubiquitination of the epidermal growth factor receptor, Genes & development, 12 (1998) 3663–3674.

[7] A.G. Batzer, D. Rotin, J.M. Urena, E.Y. Skolnik, J. Schlessinger, Hierarchy of binding sites for Grb2 and Shc on the epidermal growth factor receptor, Mol Cell Biol, 14 (1994) 5192–5201.

[8] G.A. Rodrigues, M. Falasca, Z. Zhang, S.H. Ong, J. Schlessinger, A novel positive feedback loop mediated by the docking protein Gab1 and phosphatidylinositol 3-kinase in epidermal growth factor receptor signaling, Mol Cell Biol, 20 (2000) 1448–1459.

[9] S. Sigismund, E. Argenzio, D. Tosoni, E. Cavallaro, S. Polo, P.P. Di Fiore, Clathrin-mediated internalization is essential for sustained EGFR signaling but dispensable for degradation, Dev Cell, 15 (2008) 209–219.

[10] A. Sorkin, L.K. Goh, Endocytosis and intracellular trafficking of ErbBs, Exp Cell Res, 314 (2008) 3093–3106.

[11] J. Gao, B.A. Aksoy, U. Dogrusoz, G. Dresdner, B. Gross, S.O. Sumer, Y. Sun, A. Jacobsen, R. Sinha, E. Larsson, Integrative analysis of complex cancer genomics and clinical profiles using the cBioPortal, Science signaling, 6 (2013) pl1–pl1.

[12] E. Cerami, J. Gao, U. Dogrusoz, B.E. Gross, S.O. Sumer, B.A. Aksoy, A. Jacobsen, C.J. Byrne, M.L. Heuer, E. Larsson, Y. Antipin, B. Reva, A.P. Goldberg, C. Sander, N. Schultz, The cBio cancer genomics portal: an open platform for exploring multidimensional cancer genomics data, Cancer Discov, 2 (2012) 401–404.

[13] S.V. Vasaikar, P. Straub, J. Wang, B. Zhang, LinkedOmics: analyzing multi-omics data within and across 32 cancer types, Nucleic Acids Res, 46 (2018) D956–D963.

[14] A. Subramanian, P. Tamayo, V.K. Mootha, S. Mukherjee, B.L. Ebert, M.A. Gillette, A. Paulovich, S.L. Pomeroy, T.R. Golub, E.S. Lander, Gene set enrichment analysis: a knowledge-based approach for interpreting genome-wide expression profiles, Proceedings of the National Academy of Sciences, 102 (2005) 15545–15550.

15. [15] F. Oppel, S. Shao, M. Schurmann, P. Goon, A.E. Albers, H. Sudhoff, An Effective Primary Head and Neck Squamous Cell Carcinoma In Vitro Model, Cells, 8 (2019).

[16] S. Zhou, A. Hassan, T. Kungyal, S. Tabaries, J. Luna, P.M. Siegel, A. Philip, CD109 Is a Critical Determinant of EGFR Expression and Signaling, and Tumorigenicity in Squamous Cell Carcinoma Cells, Cancers (Basel), 14 (2022).

[17] F. Ghanbari, S. Mader, A. Philip, Cholesterol as an Endogenous Ligand of ERRalpha Promotes ERRalpha-Mediated Cellular Proliferation and Metabolic Target Gene Expression in Breast Cancer Cells, Cells, 9 (2020).

[18] A.A. Bizet, K. Liu, N. Tran-Khanh, A. Saksena, J. Vorstenbosch, K.W. Finnson, M.D. Buschmann, A. Philip, The TGF-beta co-receptor, CD109, promotes internalization and degradation of TGF-beta receptors, Biochim Biophys Acta, 1813 (2011) 742–753.

[19] F. Ghanbari, A.M. Fortier, M. Park, A. Philip, Cholesterol-Induced Metabolic Reprogramming in Breast Cancer Cells Is Mediated via the ERRalpha Pathway, Cancers, 13 (2021).

[20] A. Alzahrani, Y. Chi, K.W. Finnson, M. Blati, B. Lussier, M. Kapoor, S. Roy, A. Philip, Endoglin haploinsufficiency is associated with differential regulation of extracellular matrix production during skin fibrosis and cartilage repair in mice, J Cell Commun Signal, 12 (2018) 379–388.

[21] K. Neibert, V. Gopishetty, A. Grigoryev, I. Tokarev, N. Al-Hajaj, J. Vorstenbosch, A. Philip, S. Minko, D. Maysinger, Wound-healing with mechanically robust and biodegradable hydrogel fibers loaded with silver nanoparticles, Adv Healthc Mater, 1 (2012) 621–630.

[22] L. Xu, F. Shi, Y. Wu, S. Yao, Y. Wang, X. Jiang, L. Su, X. Liu, Gasdermin E regulates the stability and activation of EGFR in human non-small cell lung cancer cells, Cell Commun Signal, 21 (2023) 83.

[23] D. Ray, A. Ahsan, A. Helman, G. Chen, A. Hegde, S.R. Gurjar, L. Zhao, H. Kiyokawa, D.G. Beer, T.S. Lawrence, M.K. Nyati, Regulation of EGFR protein stability by the HECT-type ubiquitin ligase SMURF2, Neoplasia, 13 (2011) 570–578.

[24] A. Sapmaz, A.E. Erson-Bensan, EGFR endocytosis: more than meets the eye, Oncotarget, 14 (2023) 297–301.

[25] Z. Lu, G. Jiang, P. Blume-Jensen, T. Hunter, Epidermal growth factor-induced tumor cell invasion and metastasis initiated by dephosphorylation and downregulation of focal adhesion kinase, Molecular and cellular biology, 21 (2001) 4016–4031.

[26] A.V. Vieira, C. Lamaze, S.L. Schmid, Control of EGF receptor signaling by clathrin-mediated endocytosis, Science, 274 (1996) 2086–2089.

[27] Y. Du, F. Karatekin, W.K. Wang, W. Hong, G.T.K. Boopathy, Cracking the EGFR code: Cancer biology, resistance mechanisms, and future therapeutic frontiers, Pharmacol Rev, 77 (2025) 100076.

[28] L.M. Grovdal, E. Stang, A. Sorkin, I.H. Madshus, Direct interaction of Cbl with pTyr 1045 of the EGF receptor (EGFR) is required to sort the EGFR to lysosomes for degradation, Exp Cell Res, 300 (2004) 388–395.

[29] P. Ray, K. Raghunathan, A. Ahsan, U.S. Allam, S. Shukla, V. Basrur, S. Veatch, T.S. Lawrence, M.K. Nyati, D. Ray, Ubiquitin ligase SMURF2 enhances epidermal growth factor receptor stability and tyrosine-kinase inhibitor resistance, The Journal of biological chemistry, 295 (2020) 12661–12673.

[30] P. Kavsak, R.K. Rasmussen, C.G. Causing, S. Bonni, H. Zhu, G.H. Thomsen, J. Wrana, Smad7 binds to Smurf2 to form an E3 ubiquitin ligase that targets the TGF-b receptor for degradation, Molecular cell, 6 (2000) 1365–1375.

[31] X. Lin, M. Liang, X.H. Feng, Smurf2 is a ubiquitin E3 ligase mediating proteasome-dependent degradation of Smad2 in transforming growth factor-beta signaling, The Journal of biological chemistry, 275 (2000) 36818–36822.

[32] S. Bonni, H.R. Wang, C.G. Causing, P. Kavsak, S.L. Stroschein, K. Luo, J.L. Wrana, TGF- beta induces assembly of a Smad2-Smurf2 ubiquitin ligase complex that targets SnoN for degradation, Nature cell biology, 3 (2001) 587–595.

[33] Y. Zhang, C. Chang, D.J. Gehling, A. Hemmati-Brivanlou, R. Derynck, Regulation of Smad degradation and activity by Smurf2, an E3 ubiquitin ligase, Proceedings of the National Academy of Sciences of the United States of America, 98 (2001) 974–979.

[34] L. Izzi, L. Attisano, Regulation of the TGF-b signalling pathway by ubiquitin-mediated degradation, Oncogene, 23 (2004) 2071–2078.

[35] E. Fukunaga, Y. Inoue, S. Komiya, K. Horiguchi, K. Goto, M. Saitoh, K. Miyazawa, D. Koinuma, A. Hanyu, T. Imamura, Smurf2 Induces Ubiquitin-dependent Degradation of Smurf1 to Prevent Migration of Breast Cancer Cells, J. Biol. Chem., 283 (2008) 35660–35667.

[36] P. Koganti, G. Levy-Cohen, M. Blank, Smurfs in Protein Homeostasis, Signaling, and Cancer, Frontiers in oncology, 8 (2018) 295.

[37] A. Bizet, K. Liu, N. Tran-Khanh, A. Saksena, J. Vorstenbosch, K. Finnson, M. Buschmann, A. Philip, The TGF-b co-receptor, CD109, promotes internalization and degradation of TGF-b receptors, Biochim Biophys Acta: Molecular Cell Research, 1813 (2011) 742–753.

[38] A.A. Bizet, N. Tran-Khanh, A. Saksena, K. Liu, M.D. Buschmann, A. Philip, CD109- mediated degradation of the TGF-b receptors and inhibition of TGF-b responses involve regulation of Smad7 and Smurf2 localization and function, Journal of cellular biochemistry, 113 (2012) 238– 246.

[39] J. Eom, Y. Chun, H.R. Chang, SMURF2 in Anticancer Therapy: Dual Role in Carcinogenesis and Theranostics, International journal of molecular sciences, 27 (2026).

[40] S. Hagiwara, Y. Murakumo, T. Sato, T. Shigetom, K. Mitsudo, I. Tohnai, M. Ueda, M. Takahashi, Up-regulation of CD109 expression is associated with carcinogenesis of the squamous epithelium of the oral cavity, Cancer science, 99 (2008) 1916–1923.

[41] P.O. Ozbay, T. Ekinci, S. Yigit, A. Yavuzcan, S. Uysal, F. Soylu, F. Cakalagaoglu, Investigation of prognostic significance of CD109 expression in women with vulvar squamous cell carcinoma, Onco Targets Ther, 6 (2013) 621–627.

[42] T. Sato, Y. Murakumo, S. Hagiwara, M. Jijiwa, C. Suzuki, Y. Yatabe, M. Takahashi, High-level expression of CD109 is frequently detected in lung squamous cell carcinomas, Pathology International, 57 (2007) 719–724.

[43] S. Zhu, X. Zhang, W. Liu, Z. Zhou, S. Xiong, J. Li, X. Chen, C. Peng, Ubiquitination in cancer: mechanisms and therapeutic opportunities, Cancer Commun (Lond), 45 (2025) 1128–1161.

