## Supplementary files for "A CD109-SMURF2 Axis Diverts EGFR from Degradation to Sustain Oncogenic Signaling and Promote Squamous Cell Carcinoma Invasion and Stemness"

**Supplementary methodology:**

**1. Cell sorting and validation**

In the FaDu CD109 KO experiment, single-cell sorting was conducted on the KO cell pool using FACS. Cells were sorted into CD109 KO and CD109 WT populations with the BD FACSAria Fusion system equipped with a 561 nm yellow/green laser. The sorted cells were subsequently validated by Western blotting and gene sequencing, with primers: forward (F) GCAGTGGGCAATGTTAACGG and reverse (R) TGGGGTTTCTCTGCTATACTACC

**2. Cycloheximide assay**

The cells were incubated at 37 °C in a 5% $CO_2$ for 24 hours. This was followed by overnight serum starvation, after which cycloheximide was added to the medium at a final concentration of 100 μg/mL for 2 hours. Subsequently, the cells were treated with EGF (100 ng/mL) for the specified time points, and cell lysates were collected for analysis of EGFR protein levels by immunoblot.

**3. FACS**

EGFR internalization in EV and CD109 OE A431 cells was analyzed using flow cytometry. Cells were grown for 24 hours followed by overnight serum starvation. Cells were treated with 100 ng/ml EGF for the indicated times, subsequently trypsinized and washed with PBS and staining buffer (2%FBS/1mM EDTA/PBS). Sample was incubated in blocking buffer (2% FBS/PBS) for 15 minutes then incubated with fluorophore tagged primary antibodies: anti EGFR AF488 (Abcam, ab193244) and anti CD109 PE (Invitrogen, 12-1099-42) for 20 minutes on ice in the dark. EGFR was detected using FACSAria III with blue laser (488nm) and the EGFR level was analyzed by BD FACSDiva 8.0.2 software.

**4. Immunocytochemistry (ICC)**

For the assessment of protein colocalization and EGFR internalization, SCC cells were seeded and cultured onto cover slips (170-C12MM, Ultident Scientific, Saint-Laurent, QC, Canada). Following a 10-minute treatment with EGF (100 ng/ml), cells were washed twice with PBS containing 0.2% Triton X-100 and then fixed in 4% paraformaldehyde in PBS for 10 minutes. Subsequently, the cells were blocked with 3% BSA at room temperature for 30 minutes. The cells were then incubated overnight at 4°C with fluorophore-conjugated primary antibodies: rabbit monoclonal anti-EGFR AF488 (Abcam, ab193244) and rabbit polyclonal anti-Smurf2 (G-Biosciences, ITA7689-100u-647). Nuclei were counterstained with SlowFade™ Gold Antifade Mountant with DAPI (ThermoFisher Scientific; S36938), and cover slips were mounted. Imaging was performed using a LSM780 laser scanning confocal microscope (Zeiss, White Plains, NY, USA) with a 63× oil objective.

**5. Cell invasion assay**

Corning Matrigel matrix (Corning; 356234) was coated on cellQART 24-well cell culture insert (Sterlitech; 9328012). Cells were seeded in the upper chambers with serum-free medium, while the lower chambers were filled with medium containing EGF as a chemoattractant. After a 24-hour incubation period to facilitate cell invasion through the matrix, the migrated cells were fixed with 100% methanol and subsequently stained with Giemsa. The membranes were then examined under a microscope, and images were captured to assess cell invasion.

**Supplementary data:**

**A** **B**

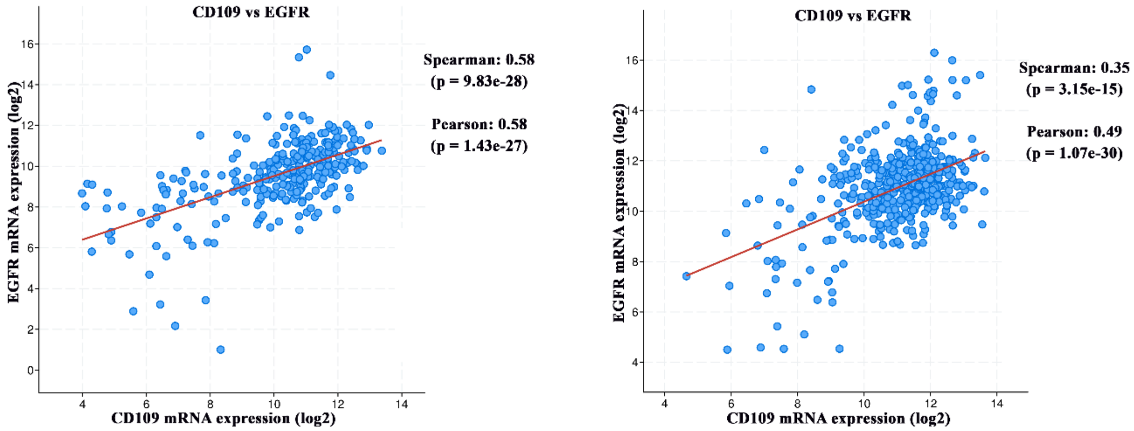


**Supplementary Figure S1. Correlation between CD109 and EGFR expression in squamous cell carcinoma datasets.** Correlation analysis of CD109 and EGFR mRNA expression in squamous cell carcinoma (SCC) patient cohorts from The Cancer Genome Atlas (TCGA) accessed through the cBioPortal database. **(A)** Pearson correlation analysis of CD109 and EGFR expression in cervical SCC patients (n = 297). **(B)** Pearson correlation analysis of CD109 and EGFR expression in lung SCC patients (n = 487). Correlation coefficients (r) and corresponding P values are indicated.

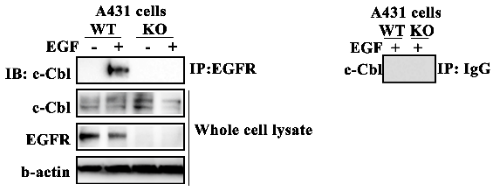


**Supplementary Figure S2. EGF stimulation enhances EGFR–c-Cbl interaction in A431 WT cells.** A431 wild-type (WT) and CD109 knockout (CD109-KO) cells were stimulated with EGF (100 ng/mL) for the indicated time points. EGFR-c-Cbl interaction was analyzed by co-immunoprecipitation. EGFR was immunoprecipitated from cell lysates and probed for c-Cbl by Western blotting. Representative Western blots showing EGFR-bound c-Cbl are presented.

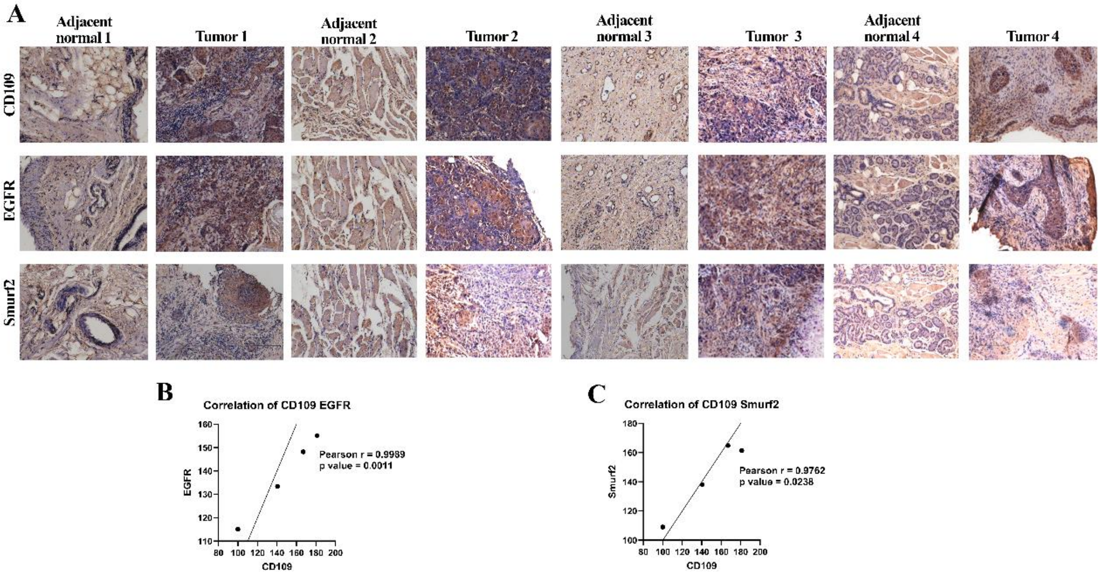


**Supplementary Figure S3. CD109 expression correlates with EGFR and Smurf2 expression in human HNSCC tissues.** (A) Representative immunohistochemical staining of CD109, EGFR, and Smurf2 in head and neck squamous cell carcinoma (HNSCC) tumor tissues and matched adjacent normal epithelium. (B-C) Correlation analysis of protein expression levels normalized to adjacent normal tissue for (B) CD109 versus EGFR expression and (C) CD109 versus Smurf2 expression in HNSCC specimens. Correlation coefficients and corresponding *P* values are indicated.

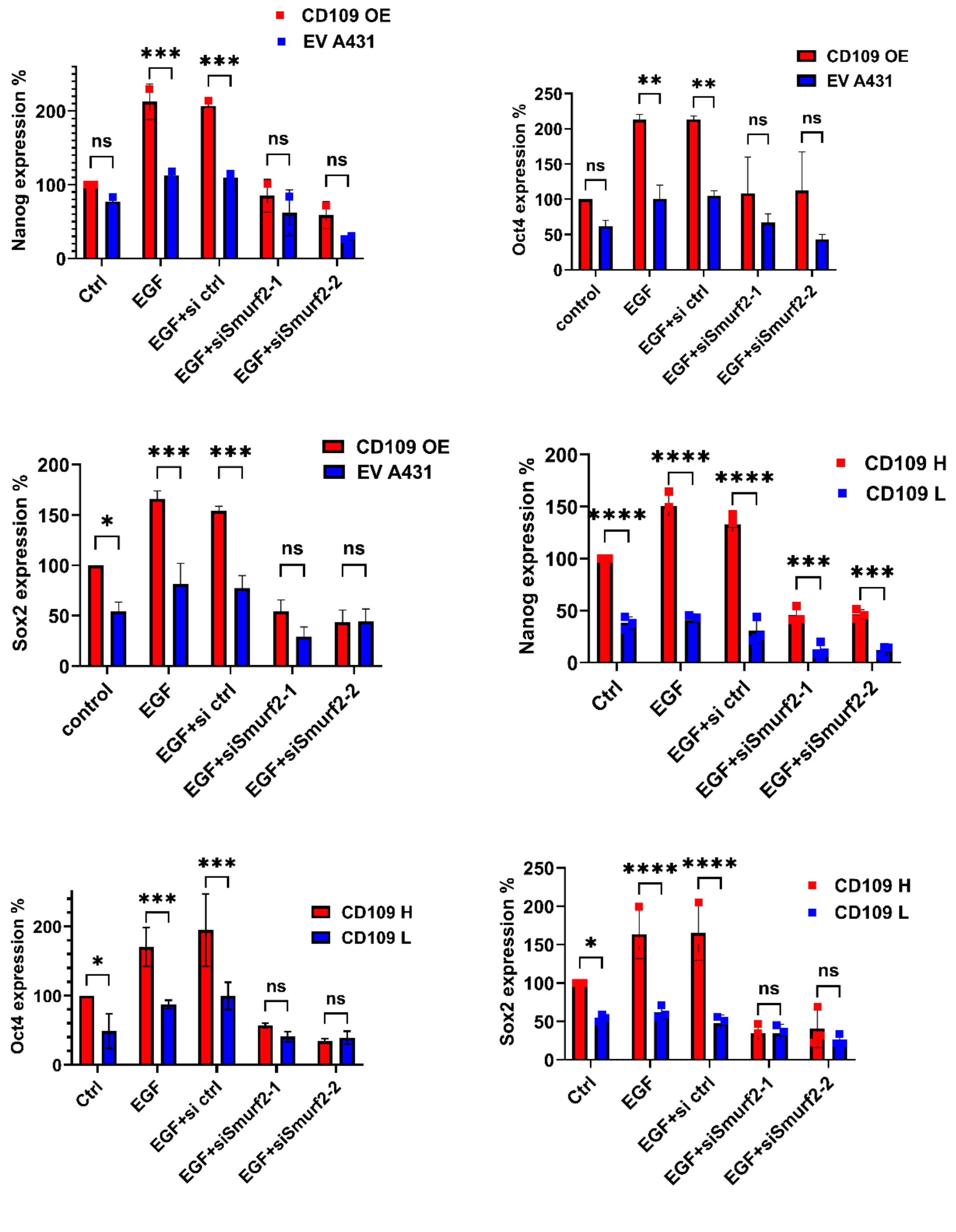


**Supplementary Figure S4. Quantitative analysis of stem cell marker expression following SMURF2 depletion. (A-C)** Densitometric quantification of stem cell marker expression in A431 empty vector (EV) and CD109-overexpressing (CD109-OE) cells following Smurf2 knockdown, from Fig. 7. Protein levels of **(A)** Nanog, **(B)** Oct4, and **(C)** Sox2 were quantified from Western blot analyses and normalized to loading control. **(D-F)** Densitometric quantification of stem cell marker expression in primary head and neck squamous cell carcinoma (HNSCC) cells stratified by CD109 expression following Smurf2 knockdown. Protein levels of **(D)** Nanog, **(E)** Oct4, and **(F)** Sox2 were quantified and normalized to loading control. Data represent mean ± SEM from at least three independent experiments. Statistical significance was determined using unpaired two-tailed Student's *t*-test. *P < 0.05; **P < 0.01; ***P < 0.001.
